# DNA condensate-based organelles for spatially regulated gene expression and protein targeting in synthetic cells

**DOI:** 10.64898/2026.05.27.728287

**Authors:** Nastasja Kaletta, Martin Matl, Petra Schwille

## Abstract

Spatial organization of gene expression is a key feature of cells but remains a major challenge in bottom-up synthetic biology. While phase-separated DNA condensates have been used to spatially confine transcription, achieving efficient recruitment of protein-coding DNA for full protein biosynthesis within these membrane-less structures has remained a major challenge. Here, we present a modular DNA nanostructure that enables tunable and highly efficient partitioning of long client DNA into condensate-based synthetic nuclei, thereby surpassing present limitations. This serves as the starting point for a multimodal DNA organelle-based system for spatially regulated gene expression in synthetic cell environments. The flow of information includes localized transcription within this synthetic nucleus, followed by translation and product release into the surrounding cytosol. Furthermore, we extend the toolkit of spatiotemporal organization by designing protein targeting of cell-cycle protein ParR within *parC*-enriched orthogonal DNA condensates. Quantitative analysis reveals a trend toward higher protein yield in the condensate-based system compared to standard PURE expression, indicating a functional advantage of spatial organization even in such minimal systems. Hence, we demonstrate how protein expression can be engineered not only by molecular composition, but by spatial architecture itself.

**Graphical abstract:** 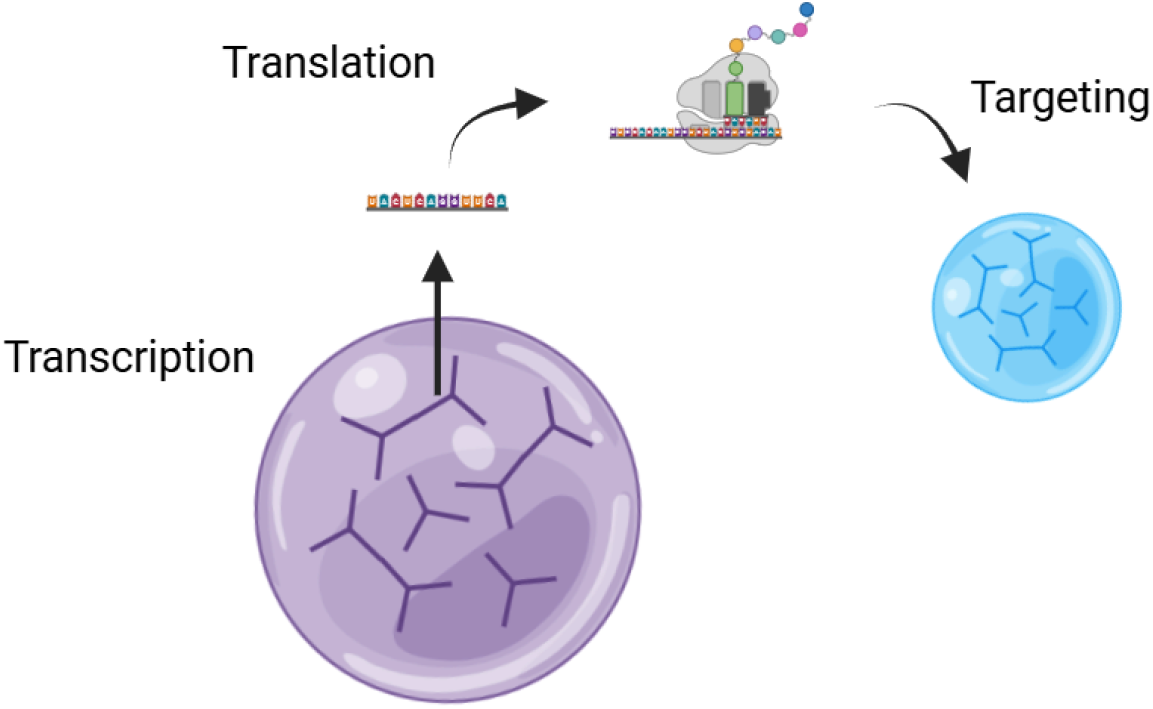

## Main

Spatial organization is a defining feature of eukaryotic cells, in which nuclear architecture, chromatin arrangement, and organelle-specific targeting coordinate transcription, translation, and protein localization to orchestrate complex cellular processes such as cell division and signal transduction [1, 2]. Furthermore, biomolecular condensates have been shown to provide an additional regulatory layer by controlling gene expression [3–5]. Recreating such architectures in cell-free or synthetic systems is a central goal in bottom-up synthetic biology and biotechnological applications [6–9]. Furthermore, the association of malfunctions in nuclear condensates with diseases such as cancer, ribosomopathy, and neurodegeneration, motivates research on elucidating the underlying mechanisms of spatial segregation [10–12].

Liquid-liquid phase separated (LLPS) condensates have emerged as a promising approach to mimic cellular compartmentalization of biochemical reactions [7]. Protein- or polymer-based condensates can locally up-concentrate specific macromolecules, for example when the molecule is modified by a specific interaction site, resulting in its enrichment [13]. PEG/dextran droplets, which are based on nonspecific interaction between DNA and the dextran phase, were used as membrane-less protocells capable of expressing protein and synthesizing building blocks for droplet self-growth [14]. Due to their lack of specificity, these LLPS systems usually fail to achieve sequence-level programmability and, therefore, controllability. Recently, a study was able to demonstrate spatial separation of transcription and translation in dextran droplets by creating a droplet-in-droplet hierarchy with protein condensates [15]. However, since the system still relies on spontaneous segregation of molecules, it does not provide the multi-step control over gene expression required to recapitulate eukaryotic organizational logic. In addition, dextran-based systems have been reported to reduce protein yield relative to bulk reactions [15]. Hence, despite successful compartmentalization, the scalability of targeted protein expression in existing systems is limited.

Programmable DNA-based condensates provide a fundamentally different solution. By carrying sequence-encoded information into the phase separation, DNA or RNA condensates enable precisely controllable, orthogonal interactions that allow selective recruitment of DNA, RNA or protein clients [16–21]. The molecular specificity these LLPS systems provide enabled the creation of artificial transcriptional condensates that not only spatially organize transcription but also allow control over downstream reactions, co-transcriptional condensation, and nuclear architectures that depend on specific oligonucleotide interactions [6, 22]. Moreover, DNA condensates have demonstrated the potential to extend these systems with additional functionalities, such as division, stimuli-responsiveness, condensate-based logic gate operations, and controllable membrane interactions inside lipid vesicles [8, 16, 20, 23]. This makes DNA condensate-based systems highly customizable and scalable.

Interestingly, gene-coding DNA hydrogels were initially reported to produce up to 300-fold higher protein levels compared to buffer solution [24]. However, in a subsequent study across multiple DNA hydrogel systems no significantly increased protein yield could be observed, calling the experimental setup of the previous work into question [25]. Nevertheless, DNA-based LLPS systems remain a promising approach for spatially organizing gene expression, as biological condensates are known to optimize the local environment of biochemical reactions and enhance reaction kinetics [7].

In particular, phase-separated DNA Y-motifs appear to be well suited as synthetic nuclei by mimicking chromatin in composition (DNA, protein), containing genetic information and compartmentalizing transcription of these genes. However, the partitioning of long DNA clients into Y-motif condensates has been limited to short sequences of approximately 202 bp due to constraints imposed by the entropic cost of confinement [26]. Importantly, protein-coding genes typically span several hundred base pairs. Hence, overcoming this limitation in client partitioning represents a key technical challenge in functionalizing DNA condensates as synthetic nuclei for synthetic biology or programmable bioreactors.

In this work, we introduce modular DNA nanostructures that achieve highly efficient partitioning of protein-coding DNA into DNA Y-motif condensates. Enabled by this advance, we establish multimodal DNA organelles that coordinate transcription within the synthetic nucleus, translation and product release into the surrounding cytosol. GFP expression was demonstrated in bulk as well as confined in water-in-oil droplets. Furthermore, we efficiently target cell-cycle protein ParR to its specific binding partner sequence *parC* in another DNA condensate compartment acting as target organelle. Quantitative analysis indicates a trend toward higher protein yield compared to standard in vitro expression, suggesting a functional advantage of spatial organization. Together, these results establish programmable DNA Y-motifs as a powerful design principle for synthetic nuclei and other functional organelles in synthetic cells. The system enables protein expression and targeting, to be controlled not only by molecular composition but by spatial orgenization of DNA condensates.

## Results and Discussion

### Design of Phase-separating DNA Nanostructures Enables Partitioning of Client DNA

To establish a modular platform for DNA nanostructure partitioning into DNA condensates as model nuclei, we initially designed a simple architecture that demonstrated basic compartmentalization of client DNA.

Our nucleus model is based on phase-separated DNA Y-motif condensates serving as the scaffold, while a second nanostructure harboring a longer DNA double strand is introduced at a lower concentration as the client molecule. This scaffold–client architecture follows a design principle conceptually similar to that described by Do et al., enabling modular recruitment without perturbing the underlying condensate structure or its biophysical properties [20]. The nanostructure for client DNA partitioning was composed of five oligonucleotides: Four of them hybridized through annealing into a four-armed DNA nanostar. Three of the arms contained palindromic sequences that facilitated transient DNA–DNA interactions. The fourth arm was extended and hybridized with the fifth oligonucleotide, which formed the double-stranded client DNA. A thymine (T_2_)-linker was introduced between the client DNA region and the nanostar arm to provide flexibility to the client DNA (Figure 1a). The scaffolding Y-motifs in our system are not only optimized to recapitulate the characteristic viscoelastic properties of membraneless organelles, but also to support efficient protein diffusion. The mesh size of 16 bp, 3-armed Y-motifs was suggested to enable the diffusion of T7 RNA Polymerase (T7 RNAP), the RNA polymerase used in this study [6, 26]. Since the partitioning of large molecules is hindered by a confinement penalty, we further extended the arm length to 40 bp. We choose 6-nucleotide overhangs to enable strong DNA–DNA interactions. In addition, we designed for two unpaired adenine bases per arm to simultaneously keep the viscosity of DNA droplets low as shown by Lee et al. [27]. Thermal annealing of newly designed oligonucleotides of scaffolding Y-motifs resulted in micrometer-sized spherical droplets. Confocal time-lapse imaging of DNA droplets following the addition of a fluorescently labelled client DNA nanostructure revealed partitioning over time (Fig 1b). Initially, the client nanostructure accumulated preferentially at the condensate interface. Intensity profiles of clients at the condensate phase transitioned from interfacial enrichment to a homogeneous flat-top distribution, consistent with equilibration and stable incorporation into the condensate interior (Fig 1c+d). This confirms successful client partitioning and transient DNA–DNA interactions within the condensates.

**Figure 1:**
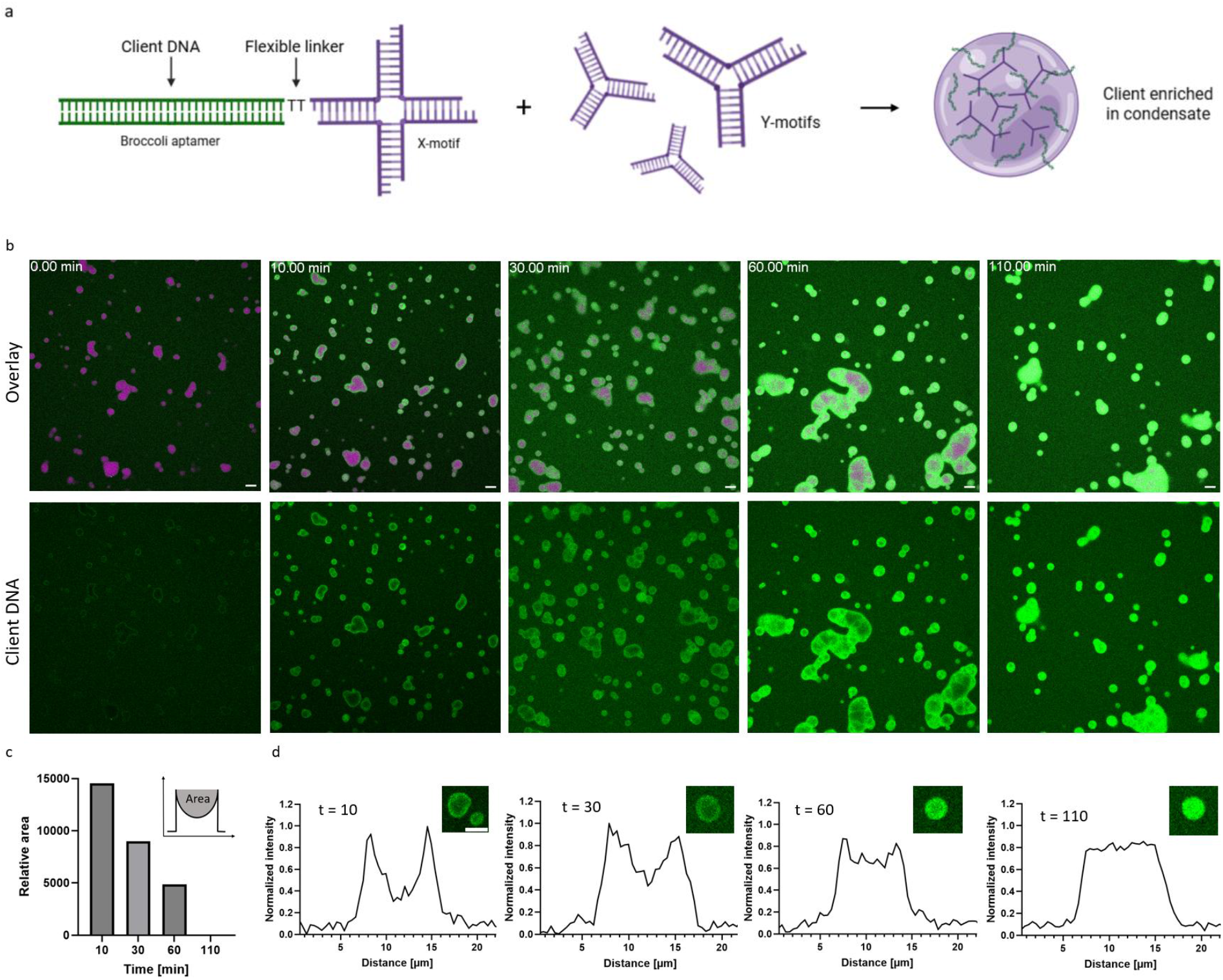
Design of phase-separating DNA nanostructures enables partitioning of client DNA A) Schematic view of DNA nanostructure for partitioning of client DNA phase separating with scaffold DNA Y-motifs B) Serial confocal images of client DNA partitioning into DNA droplets over a time course of 110 min. Green fluorescence intensity corresponds to ATTO 488-modified Y-motif DNA nanostructure for partitioning of 116 bp long client DNA. Magenta indicates ATTO 655-modified DNA Y-motifs. Shown is the green channel individually and merged images. C) Quantification of relative area representing lower client intensity at condensate interior calculated from intensity profiles in panel d). D) Mean intensity profiles of client nanostructure at condensate phase at selected time points (n = 3). Scale bars: 10 µm.

Hence, the scaffold–client DNA design we propose allows for successful partitioning of client DNA with a length of 116 bp into liquid-liquid phase separated DNA droplets.

### Broccoli Aptamer Expression Demonstrates Efficient Targeting of Transcription to the Synthetic Nucleus

The nucleus is the site of RNA transcription. Transcriptional condensates regulate gene expression by concentrating transcriptional machinery, modulating the physical environment, and selectively including or excluding molecular components [3–5].

To evaluate transcriptional activity within the DNA condensates, we employed the fluorogenic RNA aptamer Broccoli as read out. The Broccoli aptamer was encoded in the client nanostructure we partitioned into Y-motif scaffolds shown in Figure 1. Since T7 RNAP mediated transcription requires magnesium ions, DNA condensates were formed in a buffer composed of 40 mM magnesium chloride, 50 mM sodium chloride and 40 mM Tris-HCl (pH 8.0). Upon *in situ*-transcription by T7 RNAP in the presence of DFHBI-1T, a time-dependent increase in fluorescence was observed specifically within the condensates, indicating localized production of the aptamer (Fig. 2a+b, Movie 1). These findings demonstrate that DNA condensates provide a permissive and spatially confined environment for transcription.

**Figure 2:**
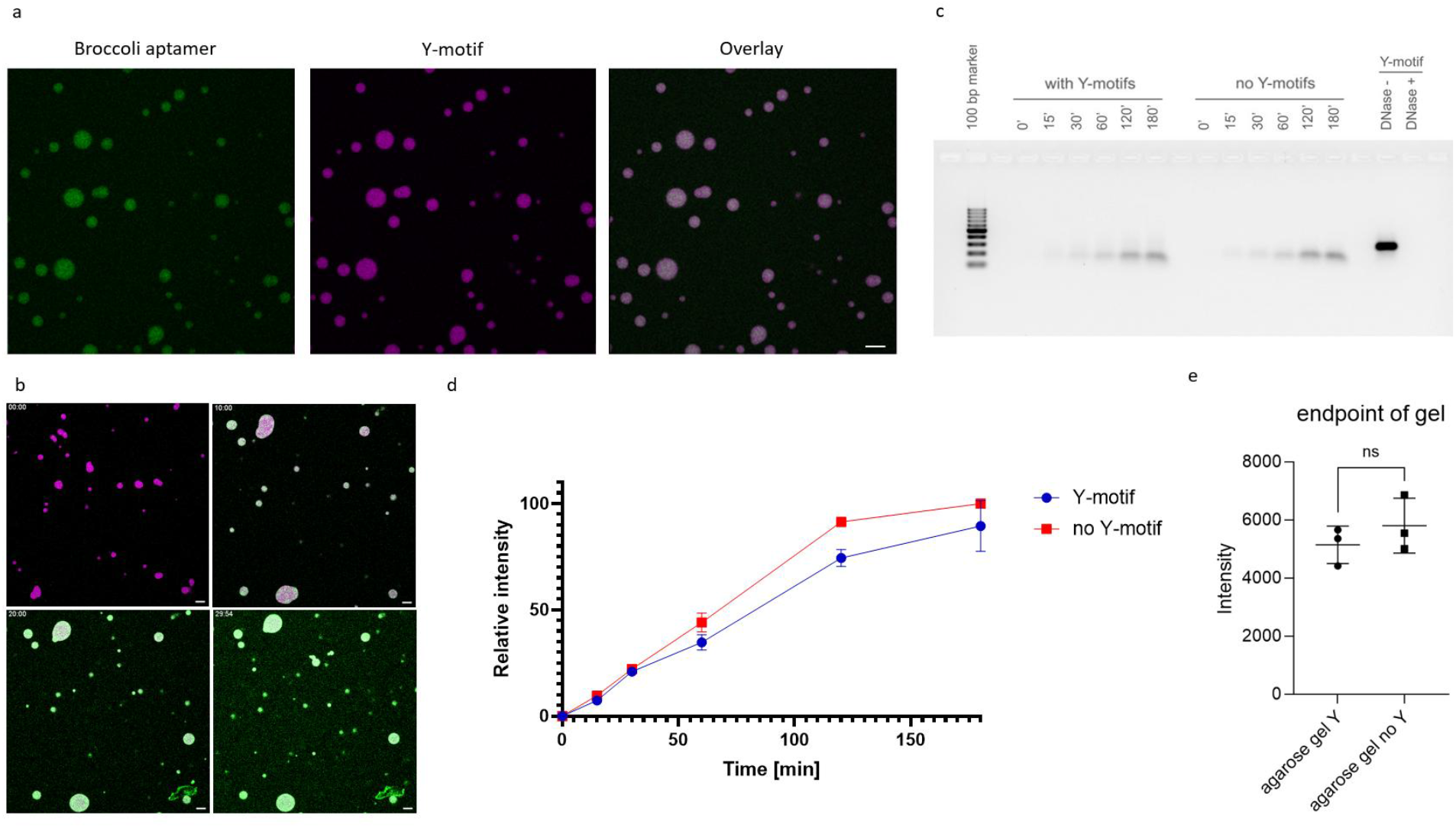
Broccoli aptamer expression demonstrates efficient targeting of transcription to the synthetic nucleus. A) Confocal images of Broccoli aptamer (green) transcribed in ATTO 655-modified DNA Y-motifs B) Serial confocal images of Broccoli aptamer (green) transcribed in ATTO 655-modified DNA Y-motifs C) 0.8 % Agarose gel of transcribed Broccoli aptamer at different time points with and without DNA Y-motifs. DNA was digested by DNase I. Gels were stained by SYBR safe. D) Quantification of transcription kinetics via densitometric analysis of agarose gels (n =3). E) Statistical analysis of end point levels of Broccoli aptamer levels via Student t-test. Data are represented as mean ± s.d.

The kinetics and transcription efficiency were quantified using agarose gel analysis (Fig 2c). Samples were treated with DNase I prior to gel electrophoresis to ensure optimal densitometric quantification of RNA bands without the DNA Y-motif band in proximity. Efficiency of DNA Y-motif digestion was confirmed (Figure 2c). Bulk kinetics of samples with and without condensates were qualitatively similar and quantitative differences in production yields were non-significant (Fig 2d+e). This is in line with observations by Kengmana et al., who did not observe significant differences in transcription rates for an RNA that was only 4 nucleotides shorter than the Broccoli aptamer when the reaction was targeted to Y-motif condensates [6].

Together, these findings confirm that the defining nuclear function of transcription can be spatially confined to our proposed DNA condensates while maintaining expression levels comparable to bulk conditions.

### Modular Design for Efficient Recruitment of Protein-coding DNA into Synthetic Nuclei

A key technical challenge in engineering synthetic nuclei is the efficient recruitment of protein-coding double-stranded DNA into DNA-based condensates. Due to their size, charge, and the entropic cost of confinement, long DNA molecules are typically excluded from dense phase-separated environments, limiting their use in synthetic compartmentalization [26].

To overcome this limitation, we first focused on optimizing the partitioning efficiency of DNA clients by tuning the architecture of the client nanostructure. Previous analyses have demonstrated that the use of longer overhangs as well as the selection of nucleotides capable of forming a higher number of hydrogen-bond interactions results in a measurable gain in Gibbs free-energy. Specifically, both contribute favourably to ΔG through enthalpic stabilization, lowering the energy cost of condensation and therefore also partitioning [16, 26]. It is likewise established that an increased number of nanostar arms can enhance the association of DNA nanostars [16]. Therefore, we hypothesised that increasing the number of arms favours partitioning of client DNA.

To test effects on partitioning by different arm numbers, we designed variants of our client nanostructure, featuring two, four and five arms. We quantified partitioning by performing confocal microscopy on fluorescently labelled versions of these nanostructures after incubation with the three-armed Y-motif scaffold (Figure 3a+b). While the design with two arms yielded only modest partitioning, the addition of a fourth arm led to a significant increase in efficiency, resulting in a ninefold higher partitioning coefficient of 42.99. Further extension to five interacting arms produced a partitioning coefficient of 64.72, corresponding to an approximately 14-fold increase in enrichment relative to the two-arm design (Fig. 3a). Nguyen et al. report accumulation of client DNA at the droplet interface for long sequences because of insufficient partitioning. Also, they observed co-phase separation of client DNA forced to partition into droplets at high concentration [26]. Despite the structural differences between the partitioning nanostructures and the three-armed Y-motif scaffold, the client DNA in our study was homogeneously distributed within the DNA droplets, and no co-phase separation was observed inside the condensates (Fig 3b).

**Figure 3:**
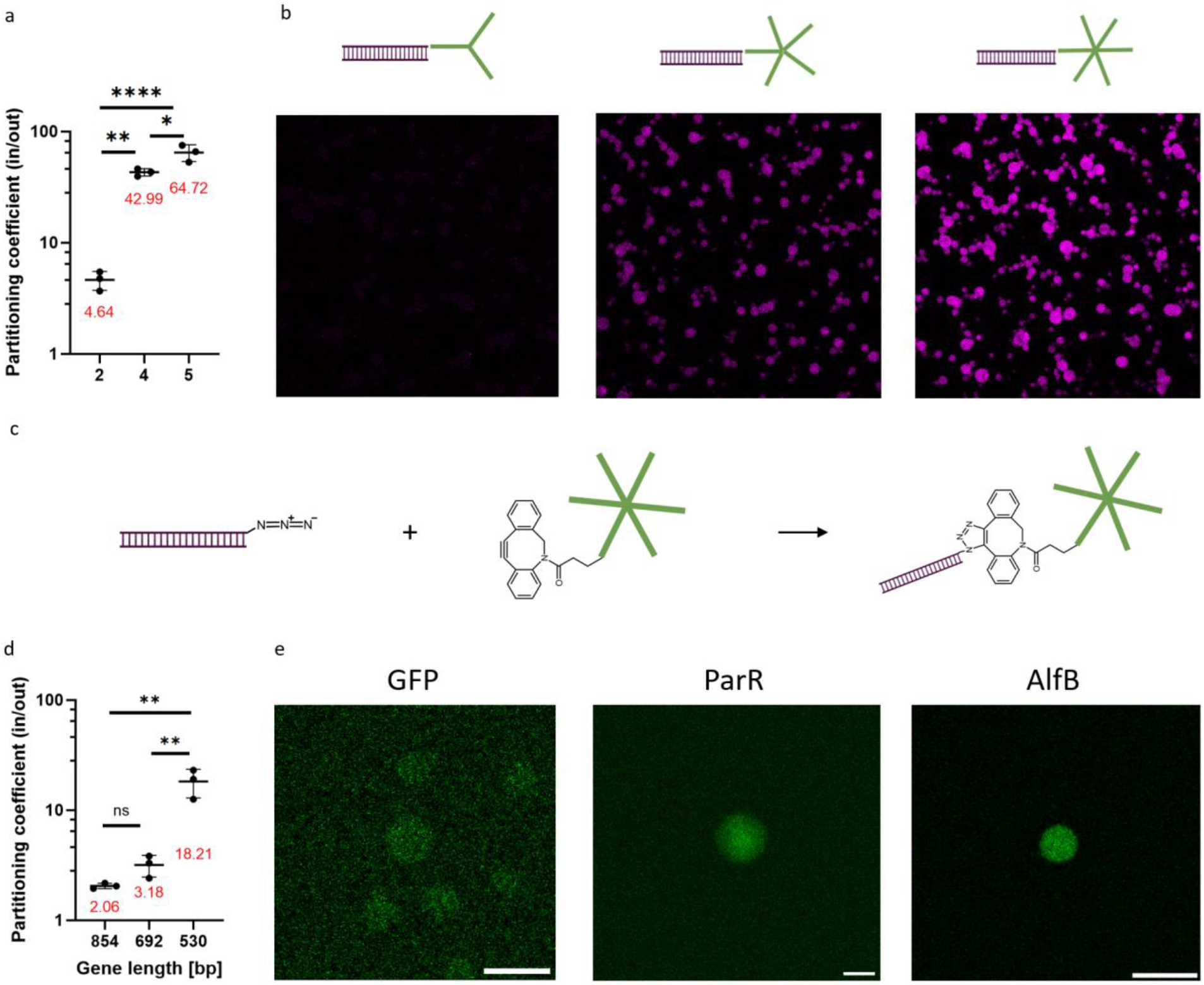
Modular Design for Efficient Recruitment of Protein-coding DNA into Synthetic Nuclei A). Statistical analysis of partitioning coefficient of client DNA with varying arm numbers in nanostructure. Data are represented as mean ± s.d. One-way ANOVA; Tukey’s multiple comparisons: *P ≤ 0.0332, **P ≤ 0.0021, ***P ≤ 0.0002, ****P ≤ 0.0001. B) Representative confocal images of 0.5 µM client DNA into DNA Y-motif scaffold. Arm number of nanostructure is indicated by schematic images. Magenta fluorescence intensity corresponds to ATTO 655-modified client. C) Schematic image of click chemistry approach to synthesize novel DNA nanostructures with gene-length DNA clients D) Statistical analysis of partitioning coefficient of client DNA with varying gene length. Data are represented as mean ± s.d. One-way ANOVA; Tukey’s multiple comparisons: *P ≤ 0.0332, **P ≤ 0.0021, ***P ≤ 0.0002, ****P ≤ 0.0001. 854 vs 692 P = 0.8980; 854 vs 530 P = 0.0016; 692 vs 530 P = 0.0024. E) Representative confocal images of 0.3 µM client DNA into DNA Y-motif scaffold. Green fluorescence intensity corresponds to ATTO 488-modified client.

Motivated by this marked improvement, we developed a protocol for functionalization of condensate-localizing nanostructures with long DNA clients capable of coding for proteins. Synthetic DNA sequences with several hundred bases are costly; therefore, we were unable to apply thermal annealing of single stranded DNA with overlaps as strategy to create the novel nanostructure with long DNA templates. To standardize the protocol for any protein and ensure efficient partitioning, we developed a click-chemistry-based strategy for assembling the nanostructure with protein-coding genes. DNA templates were amplified by conventional PCR using a primer modified with an azide group on the 5’ end. A copper-free click reaction was then performed with a DBCO-modified DNA strand to generate a defined single-stranded overhang. This overhang was designed to guide the thermal annealing of additional oligonucleotides into a six-armed DNA nanostar, with one of the arms serving as the anchor point for the client DNA double strand (Figure 3c). This approach enables site-specific functionalization of virtually any linear DNA sequence with a condensate-targeting nanostructure.

To investigate partitioning of our newly engineered nanostructures, we generated versions fused with actual gene templates coding for proteins, including GFP, ParR, and AlfB. Quantitative analysis of client-scaffold association revealed a clear dependence on DNA length. Nevertheless, even the client coding for GFP with an overall gene length of 854 bp retained the ability to partition into DNA droplets, exhibiting a partitioning coefficient of 2.06 (Fig. 3d). The spatial distribution of all novel client variants in this experiment remained homogeneous throughout the condensates, indicating efficient recruitment of gene-length clients into DNA droplets (Fig. 3e). This represents a significant improvement over previously published designs, in which effective recruitment beyond a length of 202 bp could not be demonstrated [26] and the GFP gene was instead largely excluded from the droplet phase [28].

Taken together, we established a new design strategy that enables a drastic improvement in the partitioning of client DNA. This substantial enhancement now allows us to design protein-coding client nanostructures surpassing published lengths by a factor of four [26]. Hence, we provide a general method for recruiting extended genetic programs into DNA condensate nuclei, paving the way for full protein biosynthesis in this system.

### New Client Nanostructures Enable Successful Transcription of long RNA within DNA Nuclei

Since the structure and gene length in our newly designed client nanostructures changed substantially, we evaluated whether transcription of long genes remained functional when targeted to DNA condensates.

To verify successful transcription within DNA nuclei, we designed gene templates of varying lengths, encoding different aptamers at the 5′ and 3′ ends. To fully characterize the process, the Pepper aptamer was encoded at the 5’ end and Broccoli aptamer at the 3’ end (Fig. 4a). We selected lengths of 118, 500, and 900 bp to analyze both the length-dependence and to assess whether a full transcription of our protein-coding clients is feasible inside DNA condensates. In line with the length-dependent partitioning of genes, quantification of both aptamer signals from confocal images shows a significantly higher signal for short client sequences (Figure b-e). Notably, clients with a length of 500 bp exhibit a partitioning coefficient of 73.77 for the Pepper aptamer and a partitioning coefficient of 4.60 for the Broccoli aptamer, indicating strong and complete targeting of transcription within DNA droplets, which occurs homogeneously throughout the condensates (Fig. 4). The partitioning coefficient of 1.25 for the Broccoli term of 900 bp-clients suggests that beyond this length the full transcription approaches a limiting regime, even though transcription remains targeted to DNA droplets and uniformly distributed (Fig. 4).

**Figure 4:**
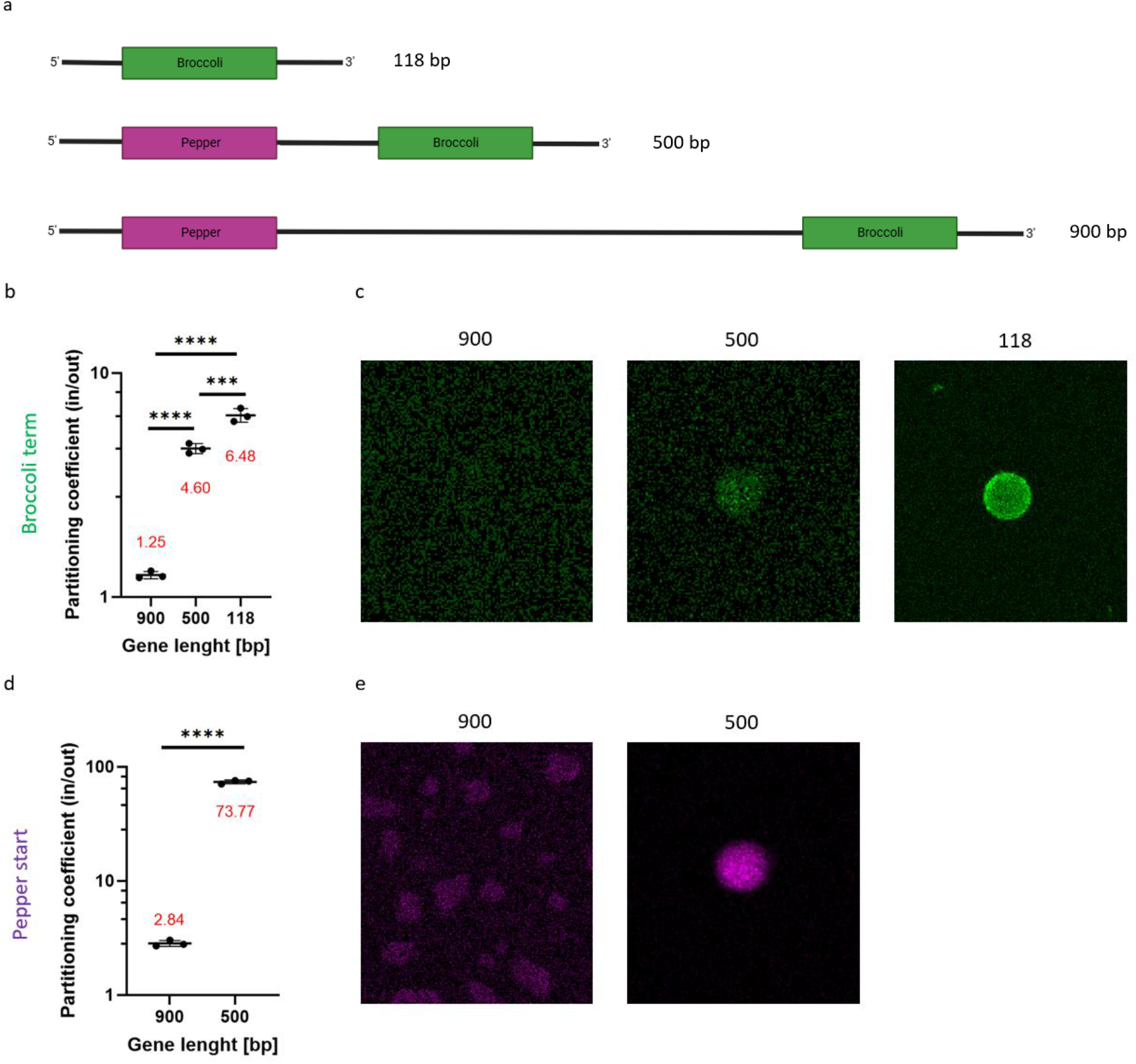
New client nanostructures enable successful transcription of long RNA within DNA nuclei. A) Schematic image of three aptamer coding gene templates of different lengths indicating location of Broccoli and Pepper aptamer. B) Statistical analysis of partitioning coefficient of transcription reaction determined by Broccoli aptamer signal. C) Representative confocal images of Broccoli aptamer signal in DNA condensates transcribed from different gene lengths (green). D) Statistical analysis of partitioning coefficient of transcription reaction determined by Pepper aptamer signal. E) Representative confocal images of Pepper aptamer signal in DNA condensates transcribed from different gene lengths (magenta). Data are represented as mean ± s.d. One-way ANOVA; Tukey’s multiple comparisons: *P ≤ 0.0332, **P ≤ 0.0021, ***P ≤ 0.0002, ****P ≤ 0.0001.

Hence, we can verify that transcription of long genes occurs successfully within the DNA droplet environment, that targeting of the reaction takes place homogeneously and can be at least for a proportion of RNA molecules fully imaged across all template-lengths within condensates.

### Condensate-Localized Genes Drive Efficient Protein Expression in Confined Cytoplasm

Having demonstrated successful targeting of protein-coding genes and their localized transcription, we characterized the complete protein biosynthesis within the DNA droplet environment. To this end, we first used green fluorescent protein (GFP) as a model protein. Coupling of transcription and translation was enabled by the PURE cell-free expression system. Time-lapse imaging of bulk reactions showed the increasing fluorescent signal of GFP in the solution surrounding DNA droplets, verifying successful translation as well as protein function (Figure 5b, Movie 2). Fluorescence imaging revealed lower GFP signal in the condensate phase, indicating product release into the surrounding solution (Supplemental Figure 1a). This could be due to charge effects as many proteins including GFP have an isoelectric point (pI) below 7, rendering them negatively charged at neutral pH, while the DNA of condensates is also negatively charged [29].

**Figure 5:**
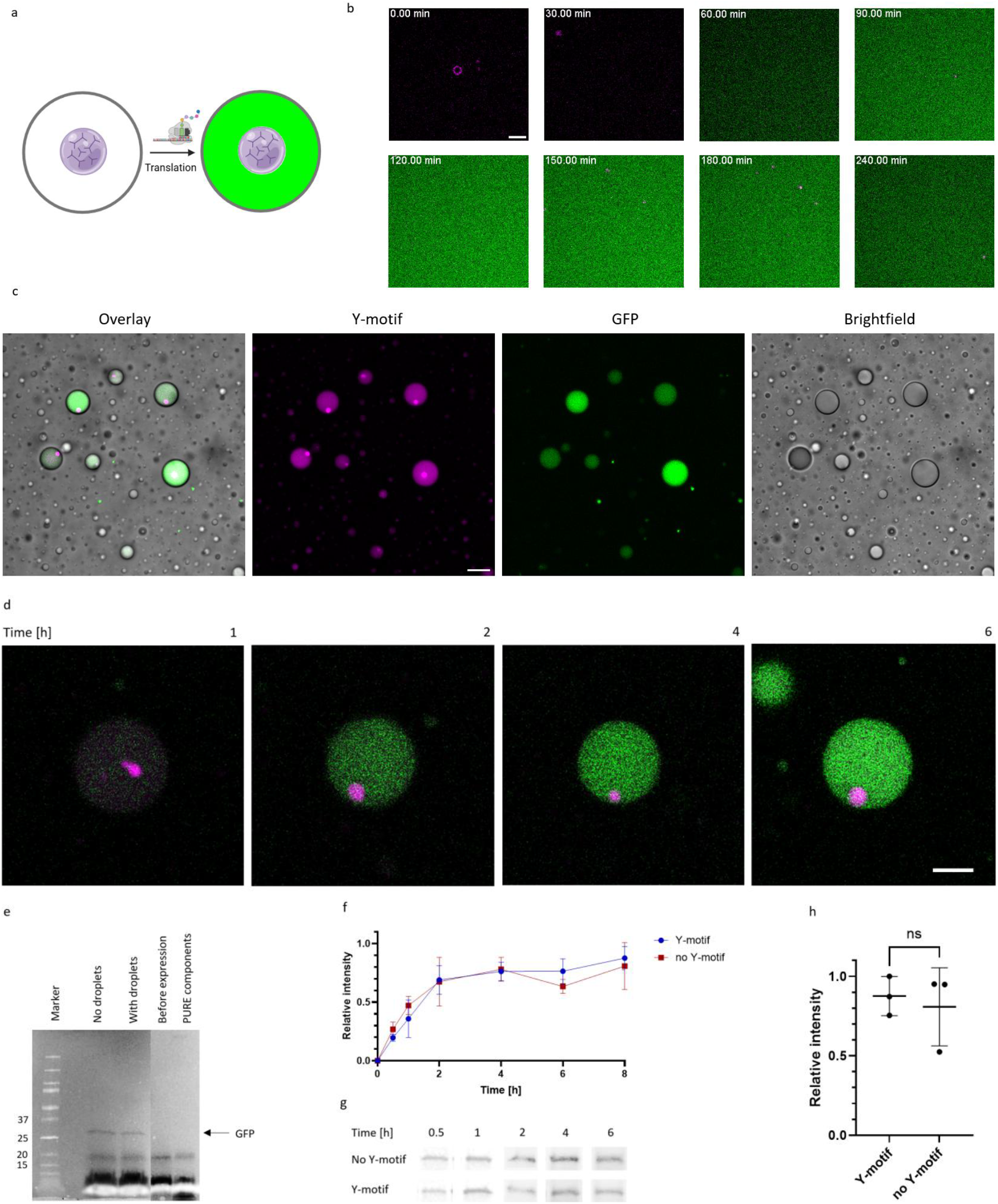
Condensate-Localized Genes Drive Efficient Protein Expression in Confined Cytoplasm. A) Schematic image of water-in-oil droplets containing a synthetic nucleus capable of expressing GFP. B) Serial confocal images of GFP expression with PUREfrex in bulk. Magenta fluorescence intensity corresponds to ATTO 655-modified Y-motif scaffold, green indicates GFP. Scale bars represent 10 μm. C) Water-in-oil droplets containing one synthetic nucleus each. GFP expression after six hours was shown. 3D Projection of water-in-oil droplets were shown in overlay, Y-motiv and GFP channel to visualize synthetic nuclei in every water-in-oil droplet. D) Confocal images of GFP expression in water-in-oil droplets containing a synthetic nucleus over time shown for several time points indicated in image. E) Denaturing SDS-PAGE of in-gel detection of GFP via FluoroTect(TM) Green(Lys) In Vitro Translation Labeling Sys. f) Quantification of GFP expression kinetics from densitometric analysis of 4-20 % SDS-PAGE with and without DNA Y-motifs. g) Representative bands corresponding to GFP at different time points from in-gel detection after expression with and without DNA Y-motifs. h) Satistical analysis of end point levels of GFP via Student t-test. Data are represented as mean ± s.d., *P ≤ 0.0332, **P ≤ 0.0021, ***P ≤ 0.0002, ****P ≤ 0.0001.

To mimic the synthetic cytosol, we demonstrated GFP production by synthetic nuclei under confinement. As a cell-sized model system, we selected water-in-oil droplets. The reaction mix was encapsulated according to our established protocol, in which preformed Y-motifs were added as final component, allowing them to form condensates under high-salt conditions within the water-in-oil droplet interiors [8]. Therefore, condensate size correlates with the water-in-oil droplet volume. To minimize interactions with the oil phase, 6 mg/ml BSA was added to the reaction mix. Water-in-oil droplets were documented after 6 hours (Figure 5c). Water-in-oil droplets containing a single cytosolic nucleus each successfully produced GFP. Hourly imaged water-in-oil droplets confirmed progressive GFP expression over time (Figure 5d). Hence, we were able to demonstrate GFP production by synthetic nuclei under cell-sized confinement.

Quantification of the GFP signal, however, revealed substantial heterogeneity between water-in-oil droplets (Figure 5c). In contrast to the bulk experiment where GFP expression reached a plateau shortly after 2 hours, signal saturation was not reached within 6 hours in the water-in-oil droplet-based system (n = 10/t) (Supplemental Figure 1b). Consistent with our observations, previous studies have reported that the GFP signal in water-in-oil droplets continues to increase for long time periods (over up to 40 hours at 30 °C)[30]. Notably, as GFP folding requires an oxidation step, the measured signal does not directly correspond to GFP expression as oxygen availability can be limited in the oil-encapsulated environment.

Therefore, we quantified expression efficiency between conditions with and without synthetic nuclei from bulk reactions. To avoid heterogeneity introduced by phase separation, we used tRNA loaded with fluorescently labeled lysine to label all *de novo* synthesized proteins prior to analysis via SDS-PAGE. In-gel detection of the labelled lysin after two hours showed a defined band at the expected migration height of GFP under conditions with and without DNA condensates (Figure 5e). Since the additional bands could be attributed to bands observed in the negative controls, we conclude that only the desired product is successfully expressed, and that no truncated or otherwise undesired by-products are generated. We next analyzed the production kinetics of GFP. As a basis for this, we performed in-gel detection of aliquots with and without DNA condensates taken after different time points. Hereby, we observed a similar kinetic profile between the samples with and without DNA droplets, with a slight trend toward increased GFP production under DNA droplet conditions (Figure 5f+g).

Reports of up to 300-fold increases in protein yield have sparked interest in the possibility of compartmentalizing DNA templates within DNA hydrogels [24]. However, recent publications employing systematic quantification across different types of hydrogels have revealed no significant differences that could be attributed to compartmentalizing of genes [25]. Since this study hypothesized that exceptionally high yields might result from excessive gene sequence concentrations within the negative control, we adhered to a concentration of 2 nM GFP gene in our experiments. After 8 hours of expression, we observed a 8 % increased protein yield in DNA droplet conditions (Figure 5h). Notably, we observed that besides GFP also the labelled negatively charged tRNA tends to be excluded from the condensate phase in confocal images (Supplemental Figure 1c). While we demonstrated successful localization of the transcription reaction within DNA condensates, many components of the translation reaction and the product (tRNA, GFP, the ribosome surface [31]) are negatively charged and seem to be rather excluded from DNA condensates. So far, the combination of synthetic condensates and protein biosynthesis has been technically challenging and associated with losses in production yield. Besides dextran droplets, a broader study across eight different liquid-liquid phase separated condensates identified one condensation system in which deGFP was generated at eightfold lower efficiency compared to the bulk solution [15, 32].

Together, this integrated control over gene localization, transcription, and functional protein output constitutes a qualitative leap beyond existing DNA-based synthetic nuclei functional in the bulk as well as under confinement, thereby providing exciting opportunities for building spatially more complex synthetic cells.

### Programmable DNA Condensates Enable Organelle-Specific Protein Targeting

Compartmentalization is a hallmark of living cells. Leveraging the unique feature of DNA Y-motifs to design orthogonality, we further extend the information stream from DNA, to RNA, to protein, to achieve efficient protein targeting into distinct receiving organelles also consisting of functionalized DNA.

To demonstrate programmable protein targeting, we established DNA-binding protein ParR as a model protein (Figure 6a). The localized ParR gene is transcribed by the synthetic nucleus. In Figure 4, we previously demonstrated the highly efficient localization of transcription of genes with similar lengths as the ParR gene (approximately 500 bp) via aptamer reporter. As a target organelle we designed a second DNA Y-motif based model organelle 2 (MO2). Targeting was enabled by enriching MO2 with ParR’s specific interaction partner, a DNA-sequence named *parC*, to achieve a spindle-assembly interaction [33, 34]. The partitioning of the *parC* sequence was achieved by client partitioning as six-armed DNA nanostructure like the gene template partitioning, which we already demonstrated in Figure 3a and b. To create individually programmable organelles, we utilized orthogonal DNA nanostars for *parC*-containing MO2, possessing overhangs with an alternative sequence. Hence, synthetic nuclei and MO2 phase separate into two distinct phases. Orthogonality of condensates under PURE conditions was confirmed by confocal imaging (Figure 6b). Accordingly, programmability of DNA Y-motifs is preserved despite the presence of magnesium ions and spermidine. Using tRNA with labeled lysine, we were able to capture the process of designed protein targeting in confocal time-lapse images. Since ParR specifically binds *parC* and no other DNA sequences, we observed an increasing fluorescence signal localized to the target organelle MO2, but not in the synthetic nuclei (Figure 6b). Quantification of the lysine signal in MO2 from time-lapse confocal images resulted in a saturation curve, which reached its plateau after approximately two hours (Figure 6c). Hence, ParR was successfully targeted to DNA condensates functionalized by interaction partner *parC*.

**Figure 6:**
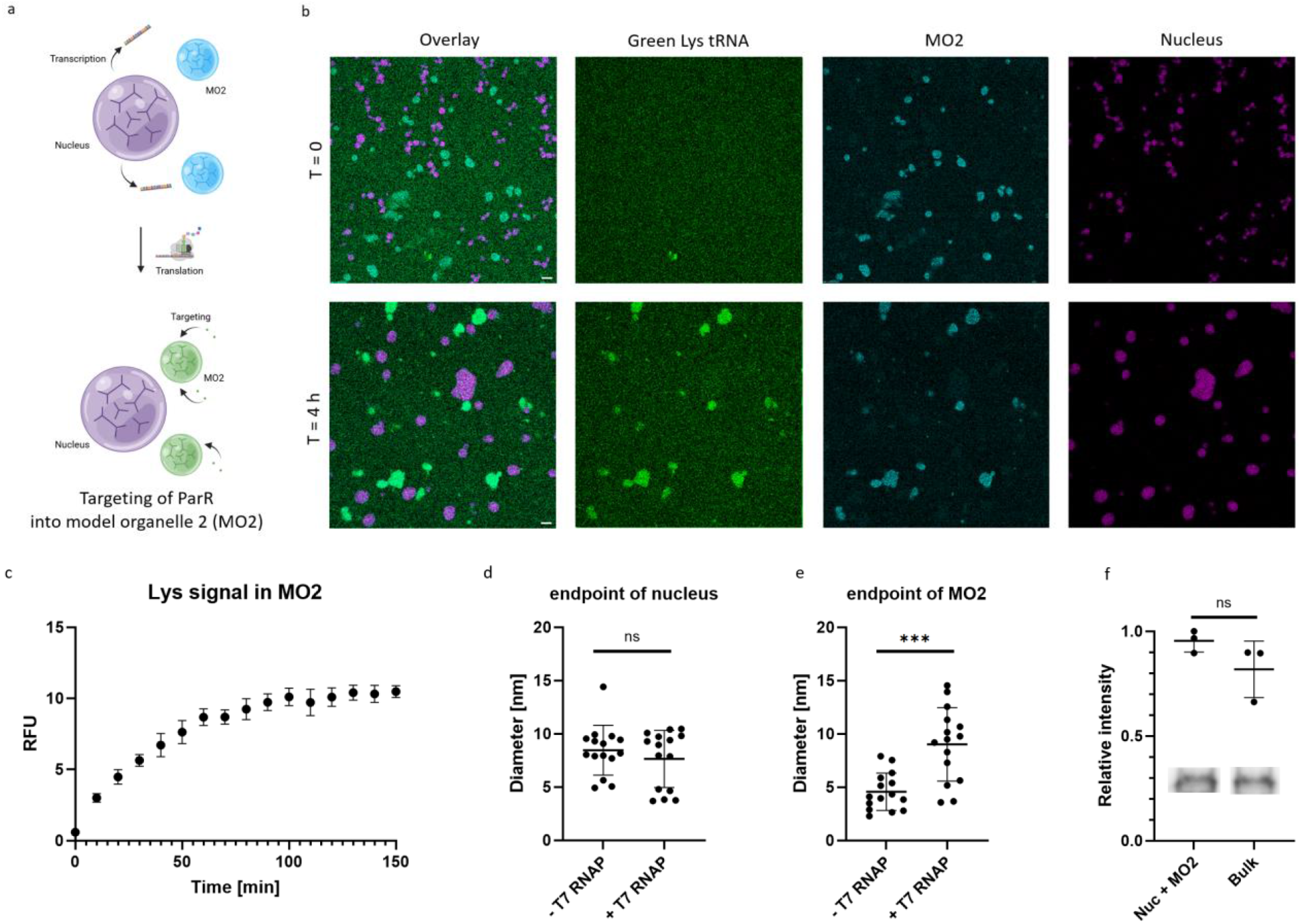
Programmable Multimodal DNA Condensates Enable Organelle-Specific Protein Targeting. A) Schematic image illustrating transcription in synthetic nucleus, translation of ParR protein in cytosol and protein targeting to parC-modified model organelle 2 (MO2) as specific binding partner. B) Confocal images before and after ParR expression with PUREfrex. Magenta fluorescence intensity corresponds to ATTO 655-modified nucleus, green indicates Green Lys for labeling of de novo synthesized ParR, cyan indicates parC-modified model organelle 2 (MO2) with ATTO 565-label. Scale bars represent 10 μm. C+D) Statistical analysis of endpoint diameters of synthetic nuclei and model organelle 2 (MO2). PURE expression without component II containing T7 RNAP was used as negative control for condensate diameter (n=15). E) Quantification of time dependent ParR levels in model organelle 2 (MO2) taken from confocal time series via FluoroTect(TM) Green(Lys) In Vitro Translation Labeling Sys for ParR labeling. F) Satistical analysis of end point levels of ParR from densitometric analysis of SDS-PAGE via in-gel detection using Student t-test. Data are represented as mean ± s.d., *P ≤ 0.0332, **P ≤ 0.0021, ***P ≤ 0.0002, ****P ≤ 0.0001. Representative bands from in-gel detection with and without DNA condensates are shown.

During the experimental procedure, we observed a change in the size ratio of the two distinct DNA condensate phases, favouring MO2 over nuclei (Figure 6d+e). Since the size of the DNA droplets changed over time due to fusion and growth events, we selected samples lacking the T7 RNAP as a control to better understand DNA organelle growth. Interestingly, there were no significant differences in nucleus size between samples that produced RNA and those that did not due to the lack of T7 RNAP (Figure 6d). Hence, the negative RNA reaction product does not seem to exert major effects on the growth of negatively charged DNA nuclei. In contrast, the binding of ParR to *parC*-functionalized MO2 resulted in a significantly larger average diameter compared to the controls, in which neither transcription nor translation of ParR occurred (Figure 6e). Hence, protein client binding facilitates DNA droplet growth. Since the target organelle is created by DNA design, our system not only facilitates precise protein targeting, but also suggests selective programmability of organelle growth.

To characterize the efficiency of protein production in the extended system, we quantified ParR yields after 8 hours via in-gel detection. We observed a 14 % higher yield under conditions with DNA condensates (Figure 6f). Although the magnitude of the improvement again falls within a range in which statistical significance is inherently difficult to demonstrate, the consistent directional trend observed during GFP expression strengthens the interpretation of a beneficial effect of compartmentation. Our results are also consistent with the findings of Tomohara et al., who reported that spatial separation of transcription and translation by PEG/Dextran-condensates resulted in a 1.2-fold increase in protein yield [15]. However, Tomohara et al. were only able to demonstrate an improvement when comparing dextran droplet conditions with coupling or uncoupling of transcription and translation. In comparison to bulk reactions, the dextran droplet system performed worse, presumably due to the presence of polymers with inhibitory effects on translation [15, 35]. In contrast to dextran droplets, our results suggest a beneficial effect on protein expression by DNA condensates when compared to standard bulk reactions. Interestingly, there is evidence suggesting that separation of transcription and translation *in vitro* can improve efficiency, as components of transcription, translation and DNA replication may exert both enhancing and inhibitory effects on these processes [36]. For example, tRNA was reported to have inhibitory effects on transcription, which appears to be excluded from the condensate phase in our system. Hence, spatial control of transcription and translation seems to also be a biochemically meaningful design strategy.

Together, our programmable DNA organelle system transcends passive, physics-driven compartmentalization by introducing orthogonality that enables true modularity. This design not only trends to increase protein yield, but also establishes a genetically addressable, multi-step expression pipeline capturing essential aspects of eukaryotic organizational logic.

## Conclusion

In this work, we establish controllable phase-separated DNA Y-motifs as a modular platform for spatial organization of cellular processes, such as gene expression, in bottom-up synthetic systems. By overcoming the key technical challenge of efficiently recruiting protein-coding DNA into phase-separated compartments, we enable full protein biosynthesis from condensate-localized genes and thereby close a critical gap between transcriptional localization and functional protein expression. By extending our system to include protein targeting, we demonstrate a comprehensive stream of information from DNA to RNA, to protein, and finally to the target organelle.

A key advance of our approach is the ability to efficiently recruit long protein-coding DNA clients into phase-separated droplets, overcoming a central limitation of existing DNA condensate-based systems [6, 22, 26]. This integrated control over gene localization, transcription, and functional protein output recapitulates characteristics of eukaryotic protein synthesis. In literature, interferences between transcription and translation, as well as DNA replication, have been observed in vitro, suggesting the spatial segregation of these biochemical processes as a promising design strategy [36]. For example, the exclusion of tRNA, which is reported to exert inhibitory effects on transcription, represents an intrinsic property of Y-motif droplets favorable for applications in synthetic biology and biotechnology.

Previous work has demonstrated that phase-separated PEG/dextran droplets can spatially organize gene expression components and support segregation of transcription and translation reactions. However, these systems rely on largely passive and nonspecific physicochemical interactions, which can lead to unintended sequestration effects of not only DNA but also RNA, polymers and proteins [14, 15, 37–40]. Additionally, in existing dextran nuclei protein yield was reduced compared to bulk reactions [15]. In contrast, our DNA-based condensates are built from sequence-defined, orthogonal interactions that decouple condensate formation from functional recruitment [16]. This programmability allows separate control over genome recruitment, transcriptional confinement, translational access, and protein targeting. As a result, our system trends toward enhancing protein expression relative to bulk conditions, highlighting that functional compartmentalization requires not only phase separation but also molecular orthogonality and designable interactions. Together, these findings emphasize that DNA-programmable condensates provide a fundamentally different and functionally advantageous framework compared to non-specific polymer-based droplets such as dextran.

The orthogonal targeting of the cell-cycle-associated protein ParR to *parC*-enriched DNA organelles highlights the biological potential of our platform. Unlike previous systems relying on artificial reporters, ParR represents a native DNA-binding protein whose localization remains effective under transcription–translation conditions.

Finally, the coexistence of multiple, compositionally distinct DNA-based condensates demonstrates that higher-order organization can be achieved through DNA design introducing multimodularity to the system. Targeted, specific compartmentalization mirrors key principles of cellular organization and underscores the potential of programmable phase separation as a key foundation for synthetic cell engineering. Besides orthogonality, protein expression from synthetic nuclei in water-in-oil droplets demonstrates even further hierarchical organization, hinting at even greater scalability of our system. While many modules in synthetic biology are traditionally lipid vesicle-based, GFP expression was reported to be readily increased in water-in-oil droplets, highlighting spatial confinement in general as attractive strategy for synthetic biology and biotechnology [30]. Furthermore, DNA condensates have demonstrated the potential to extend the system with additional functionalities, such as division, stimuli-responsiveness, condensate-based logic gate operations, and controllable membrane interactions inside lipid vesicles [8, 16, 20, 23]. In particular, the ability to program division via the orthogonality of Y-motifs represents an advantage over dextran droplets applicable in future work [14].

In summary, our results show that DNA-programmable condensates may serve as faithful mimicries of synthetic nuclei in which genetic information can be spatially localized, thereby providing the means for spatiotemporal engineering of gene expression. This may not only increase overall protein yield but also establishes a versatile platform for constructing synthetic cells with increasing organizational complexity and functional integration.

## Methods

### Annealing of DNA droplets and DNA nanostructures

Oligonucleotides were purchased from IDT (USA, purification of unmodified oligonucleotides: page, purification of modified oligonucleotides: HPLC Purification), dissolved at 100 µM in milliQ water and stored at −20 °C. DNA strand sequences are provided in supplemental tables S1, 3-4. Equimolar ratios of strands were mixed in an Eppendorf PCR tube in H_2_0 for scaffolding Y-motifs or at 12.5 µM in buffer for client nanostructures (50 mM NaCl, 40 mM Tris pH 8.0, 40 mM MgCl_2_). Strands were annealed on a Thermal Cycler (Bio-Rad, CA, USA) heated to 85 °C and cooled down to 23 °C at a rate of −1° per minute.

### Molecular cloning

T7 RNAP constructs were designed and cloned via seamless cloning. The plasmid was from Addgene https://www.addgene.org/60717/. In brief, vectors and inserts were amplified using indicated primers (Table S2) via PCR (Phusion Polymerase (ThermoFisher Scientific Cat. No. F530L)). Template was digested via DpnI (NEB) at 37 °C for 30 min. PCR products were loaded and visualized on agarose gel electrophoresis (agarose 0.8 %, TAE, 120 V, 40 min). Bands were cut-out and purified using a QIAGEN gel extraction kit (Cat. No. 28704). Following PCR of inserts, vector and insert were assembled by seamless cloning using the ThermoFisher Scientific /Invitrogen GeneArt− Seamless Cloning and Assembly Enzyme Mix (A14606). Plasmids were transformed into chemically competent OneShot Top10-cells (Cat. No. C404006). Colonies were grown on LB-Kanamycin plates at 37 °C. Clones were picked and grown overnight in 5 ml LB-Kanamycin media. Plasmids were purified via Miniprep kit (Cat. No.27104) and sequenced.

### T7 RNAP expression and purification

T7 RNA polymerase was expressed as a cleavable N-terminal 6×His-tagged protein in *E. coli* BL21 cells from a T5 lac–inducible plasmid. Cells were grown in LB medium supplemented with kanamycin and induced at an optical density of ~1.0 with 1 mM IPTG, followed by expression at 18 °C for 16 h. Cells were harvested by centrifugation, resuspended in ice-cold lysis buffer (50 mM Na_2_HPO_4_, 300 mM NaCl, 10 mM imidazole, 1 mM DTT, 1x cOmplete Mini EDTA-free protease inhibitors (Roche), and 3 mg/ml lysozyme), and lysed by sonication. Insoluble material was removed by centrifugation at 20,000 × g for 45 min at 4 °C. The clarified lysate was incubated with Ni-NTA agarose for 15 min at 4 °C and loaded onto a gravity column. The resin was washed sequentially with 10 column volumes of wash buffer containing 10 mM imidazole, 10 column volumes containing 20 mM imidazole, and 5 column volumes containing 30 mM imidazole. Bound protein was eluted with 2 column volumes each of wash buffer containing 50, 100, 150, and 200 mM imidazole. Elution fractions containing T7 RNA polymerase were pooled and dialyzed overnight at 4 °C in the presence of His-tagged TEV protease (1:30 protease-to-protein ratio) to remove the affinity tag. The dialyzed sample was subjected to reverse Ni-NTA chromatography by incubation with Ni-NTA agarose for 30 min at 4 °C. The resin was loaded onto a gravity column and washed sequentially with 2 column volumes of buffer containing 0, 5, and 10 mM imidazole, followed by elution with 500 mM imidazole. Untagged T7 RNA polymerase present in the flow-through and low-imidazole wash fractions was pooled, concentrated using 30 kDa molecular weight cut-off centrifugal concentrators, and stored at −80 °C as single-use aliquots.

### Transcription in DNA droplets

2 µM scaffolding Y-motifs, 0.2 µM client nanostructure, 2 mM NTPs, 5 mM DTT, and 30 µM DFHBI-1T (and HBC620, GLP Bio) were mixed in transcription buffer (50 mM NaCl, 40 mM Tris pH 8.0, 40 mM MgCl_2_). Solution was incubated for 30 minutes for condensate-formation in a 96-Well Flat-Bottom Microplate (SensoPlate, Greiner Bio-One GmbH, Kremsmuenster, Austria) passivated with 10 mg/ml BSA for 10 minutes. Reaction was induced by adding an enzyme mix providing a final concentration of 0.2 µM T7 RNAP, 1 µM RiboLock RNase Inhibitor (ThermoScientific) and inorganic pyrophosphatase (NEB). Transcription was monitored via confocal imaging or gel electrophoresis (agarose 1.5 %, TBE, 100 V, 90 min). If DNase digestion was needed, DNase I (NEB) was used and added to samples at a ratio of 2 µl DNase for 10 µl reaction and incubated for 30 minutes at 37 °C. DNase was inactivated by adding 50 mM EDTA and incubating at 75 °C for 10 minutes.

### Production of protein-coding client nanostructures via click chemistry

Gene templates were purchased from IDT as gBlocks (Table S5). Templates were amplified via PCR using an azide-modified primer. Click reaction was performed at 37 °C for 3 days by adding a DBCO-modified oligonucleotide able to hybridize with a star shaped nanostructure harboring single stranded overhangs. Success of click reaction was confirmed by agarose gel electrophoresis (agarose 2 %, TAE, 80 V, 120 min) showing a shift in DNA band. DNA was purified and concentrated via InnuPREP DNA/RNA Mini Kit (Analytik Jena AG, Germany). DNA concentration was measured via nanodrop and equimolar strands for client nanostructure added in buffer (50 mM NaCl, 40 mM Tris pH 8.0, 40 mM MgCl_2_). Nanostructures were formed via thermal annealing as described.

### PURE expression in DNA droplets

Cell-free expression was performed using PUREfrex 2.0 (GeneFrontier, Chiba, Japan) following the supplier’s instruction with minor changes. PURE solutions, 5 µM Y-Motif scaffold, and 2 nM gene template as client nanostructure were mixed without PURE solution I containing T7 RNAP. The solution was incubated at 37 °C to allow DNA condensate-formation. 9.5 µl reaction mix was transferred on a BSA passivated coverslip (Menzel Glasses, # 1.5, 22 mm × 22 mm, Thermo Fisher Scientific) with an imaging spacer (Grace Bio-Labs SecureSeal imaging spacers, 1 well, 9 mm × 0.12 mm, Sigma–Aldrich). Reaction was induced by adding PURE solution I and reaction chambers sealed by a second cover slip. For labeling of de novo synthesized protein 0.3 µl tRNA loaded with labelled lysine (FluoroTect(TM) Green(Lys) In Vitro Translation Labeling Sys) was added to reaction mix. Labelled proteins were identified via confocal microscopy or in-gel detection via SDS-PAGE. For protein targering experiments 0.8 µM *parC* client nanostructure and orthogonal Y-motif scaffolds were added.

### Water-in-oil droplets

For imaging in water-in oil droplets 6 mg/ml BSA was additionally added to PURE reaction mix. To wash out residual molecules, lyophilized BSA (Sigma–Aldrich, catalog number: A6003) was dissolved in buffer (50 mM Tris-HCl, pH7.5, 150 mM KCl, 5 mM MgCl_2_). The buffer was exchanged with fresh buffer after centrifugation in Amicon Ultra-0.5 centrifugal filter unit 50 kDa (Merck KGaA, Darmstadt, Germany) for three times. Droplet formation was achieved by thoroughly vortexing 5 µl PURE reaction mix in 200 µl silicone oil (Merck KGaA, Darmstadt, Germany).

### Microscopy

Microscopy images were taken on a Zeiss LSM780 confocal laser scanning microscope using a Zeiss C-Apochromat × 40/1.20 water-immersion objective or a Plan-Apochromat 20x/0.80 air objective (Carl Zeiss). A pinhole size of 2.6–4 Airy units for the channels 512 × 512-pixel resolution and a pixel dwell time of 1.27 μs was used. Z-stacks with 0.2 μm intervals were obtained for three-dimensional imaging. Microscopy images were processed via NIH Fiji ImageJ and contrast and brightness adjustments applied uniformly to the entire image field.

### Statistics

Statistical significance between conditions with and without DNA condensates were determined via one-way analysis of variance (ANOVA) with Tukey’s multiple comparisons. Statistical analysis was calculated via GraphPad Prism version 8. 4.. Statistical significance is indicated by: ****P ≤ 0.0001; ***P ≤ 0.001; **P ≤ 0.01; *P ≤ 0.05; NS (not significant) P > 0.05.

### ChatGPT for Grammar Verification and Translation

AI-based language model ChatGPT developed by OpenAI was used for language optimization and assistance with phrasing in this manuscript.

## Acknowledgements

The authors would like to thank Dr. Jan-Hagen Krohn for insightful discussions. The authors thank Michaela Schaper and Kerstin Röhrl for their support with cloning experiments, PCR and protein purification. N.K. acknowledges the International Max Planck Research School for Molecules of Life (IMPRS-ML) for support. N.K. and P.S. received funding by the European Research Council (ERC Synergy Grant MetaDivide, no. 101167181). We also acknowledge the support of the Center for Nanoscience (CeNS), Munich. Schematic images were created with BioRender.com.

## Contributions

P.S. and N.K. conceptualized the work. N.K. designed the experiments, performed the experiments, analyzed the data, and wrote the first draft of the manuscript. M. M. contributed data for selected figures. P.S. supervised the research. All authors reviewed and approved the manuscript.

