## Supplemental material for "DNA condensate-based organelles for spatially regulated gene expression and protein targeting in synthetic cells"

Nastasja Kaletta (1), Martin Matl (2, 3), Petra Schwille (1)

1 Department of Cellular and Molecular Biophysics, Max Planck Institute of Biochemistry, Am Klopferspitz 18, 82152 Martinsried (Germany)
2 Institute of Molecular Biotechnology of the Austrian Academy of Sciences (IMBA), Vienna BioCenter (VBC), Dr. Bohr-Gasse 3, 1030 Vienna, Austria
3 Vienna BioCenter PhD Program, Doctoral School of the University of Vienna, Medical University of Vienna, Vienna, Austria

Table 1: Oligonucleotide sequences of Y-motif scaffold and client DNA nanostructure for partitioning of Broccoli template. Y-12, X-3 and X-4 form scaffolds. Broccoli_X-1, Broccoli_X-1_rev_complement, X-3 and X-4 form client

| Name | Sequence (5‘ - 3‘) |
| --- | --- |
| Broccoli_X-1 | taatacgactcactataggGGTGCCTTATTCCGGACGCCGGGCCCGAATGCTGCTACGGCAGTCGAAGACAACATCGCGCCCTTCGGAGGCACCctagcataaccccttggggcctTTCAGTGAGGACGGAAGTGAAGGAACTCTCCGCGTCTCCGCGTTGTCGTAGCGTCGTAGCATCGCACCGACAAAGCGAACACGT |
| Broccoli_X-1_rev_complement | aggccccaaggggttatgctagGGTGCCTCCGAAGGGCGCGATGTTGTCTTCGACTGCCGTAGCAGCATTCGGGCCCGGCGTCCGGAATAAGGCACCcctatagtgagtcgtatta |
| X-2 | GCTAGCAAACGTGTTCGCTTTGTCGGTGCGATGCTACGACGCTACGACTTCTGATGACTTCCGTCCTCACTGGTGCTGGCATACCTGACT |
| X-3 | GCTAGCAAAGTCAGGTATGCCAGCACCAGTGAGGACGGAAGTCATCAGTTCGAGTGAGCATCTAGTGCTGTCAGACGATGCGTGCTGAGT |
| X-4 | GCTAGCAAACTCAGCACGCATCGTCTGACAGCACTAGATGCTCACTCGTTCGCGGAGACGCGGAGAGTTCCTTCACTTCCGTCCTCACTG |
| Y-12 | GCTAGCAACAGTGAGGACGGAAGTGAAGGAACTCTCCGCGTCTCCGCGTTCTGATGACTTCCGTCCTCACTGGTGCTGGCATACCTGACT |
| Y-12_ATTO633 | [ATTO633]- GCTAGCAACAGTGAGGACGGAAGTGAAGGAACTCTCCGCGTCTCCGCGTTCTGATGACTTCCGTCCTCACTGGTGCTGGCATACCTGACT |

Table 2: Primers for T7 RNAP cloning. Forward (f) and reverse (r) primers for pCA24N vector and T7 RNAP insert.

| Name | Sequence (5‘ - 3‘) |
| --- | --- |
| pCA24N_f | ggattggaagtacaggttttcatggtgatggtgatggtgaga |
| pCA24N_r | taaggcctatgcggccgc |
| T7RNAP_i_f | gaaaacctgtacttccaatccatgaacacgattaacatcgct |
| T7RNAP_i_r | gcggccgcataggccttacgcgaacgcgaagtc |

Table 3: Oligos used to form client nanostructures with two (Y), four (Five) and five (Six) arms. ParC-Y-1_6, ParC_rev_compl and Y-2_6 participated in all nanostructures. To form Model organelle 2 (MO2) exchange overhangs (GCTAGC) with orthogonal overhangs (CTCGAG).

| Name | Sequence (5‘ - 3‘) |
| --- | --- |
| ParC-Y-1_6 | GCTAGCGGTGTTTTTTTGGTGTGTGTTTGGGTATGTTTTGGGTTTTAAGTGGGTTTGTTTGtcaagtttaccccatttcaaccatcaatcaatgattattTGTCTTGTTTTGGTGTTTTATTGGGTTGGGTATGGGTTGTTTTTGGGTTTTGTTTGCTAGCTTGCTAGC CAGTGAGGACGGAAGTTTGTCGTAGCATCGCACC |
| ParC_rev_compl | GCTAGCAAACAAAACCCAAAAACAACCCATACCCAACCCAATAAAACACCAAAACAAGACAaataatcattgattgatggttgaaatggggtaaacttgaCAAACAAACCCACTTAAAACCCAAAACATACCCAAACACACACCAAAAAAACACCGCTAGC |
| Y-2_6 | GCTAGC CAACCACGCCTGTCCATTACTTCCGTCCTCACTG |
| Y-3_6 | GCTAGCGGTGCGATGCTACGACTTTGGACAGGCGTGGTTG |
| Six-3_6 | GCTAGC CCATGGTCCCAAGTGATTTGGACAGGCGTGGTTG |
| Six-4_6 | GCTAGC CGGCGCTGTAAATTTGTTTCACTTGGGACCATGG |
| Six-5_6 | GCTAGC CAGACGTCACTCTCCATTCAAATTTACAGCGCCG |
| Six-6_6 | GCTAGC GGTGCGATGCTACGACTTTGGAGAGTGACGTCTG |
| Five-3_6 | GCTAGC GCTGGACTAACGGAACTTTGGACAGGCGTGGTTG |
| Five-4_6 | GCTAGC GGTGCGATGCTACGACTTTCAGGTATGCCAGCAC |
| Five-5_6 | GCTAGC GTGCTGGCATACCTGATTGTTCCGTTAGTCCAGC |

Table 4: Primers and oligos to synthesize client nanostructure via click chemistry approach. PURE primers to amplify PURE template. Amplify with azide-modified primer for click chemistry with DBCO-Client_1.1.

| Name | Sequence (5‘ - 3‘) |
| --- | --- |
| PURE_f | CCCGCGAAATTAATACGACTCAC |
| PURE_r | CAAAAAACCCCTCAAGACCCGT |
| ATTO488-PURE_f | [ATTO 488]-CCCGCGAAATTAATACGACTCAC |
| Azide_PURE_r | [Azide]-CAAAAAACCCCTCAAGACCCGT |
| DBCO-Client_1.1 | [5DBCON]-TTCAGTGAGGACGGAAGTGAAG |
| Client_1.2 | AACTCTCCGCGTCTCCGCGTTGTCGTAGCGTCGTAGCATCGCACCGACAAAGCGAACACGT |
| Client_2 | GCTAGCAAACGTGTTCGCTTTGTCGGTGCGATGCTACGACGCTACGACTTAGAGTACGTGCGAAGCCGTCGTCGGTGCAGACCTAGTAGC |
| Client_3 | GCTAGCAAGCTACTAGGTCTGCACCGACGACGGCTTCGCACGTACTCTTTACGCCTAGTAGAGTGATGAGAGCCATGAGTGCCGCGCTAT |
| Client_4 | GCTAGCAAATAGCGCGGCACTCATGGCTCTCATCACTCTACTAGGCGTTTCTGATGACTTCCGTCCTCACTGGTGCTGGCATACCTGACT |
| Client_5 | GCTAGCAAAGTCAGGTATGCCAGCACCAGTGAGGACGGAAGTCATCAGTTCGAGTGAGCATCTAGTGCTGTCAGACGATGCGTGCTGAGT |
| Client_6 | GCTAGCAAACTCAGCACGCATCGTCTGACAGCACTAGATGCTCACTCGTTCGCGGAGACGCGGAGAGTTCCTTCACTTCCGTCCTCACTG |

Table 5: Genes used for click chemistry approach

| Name | Sequence (5‘ - 3‘) |
| --- | --- |
| GFP | cccgcgaaattaatacgactcactatagggAGACCACAACGGTTTccctctagaaataattttgtttaactttaagaaggagatataccATGAGTAAAGGAGAAGAATTATTTACTGGAGTTGTCCCAATTCTTGTTGAATTAGATGGTGATGTTAATGGGCACAAATTTTCTGTCCGTGGAGAGGGTGAAGGTGATGCAACAAACGGAAAACTTACCCTTAAATTTATTTGCACTACTGGAAAACTACCTGTTCCATGGCCAACACTTGTCACTACTTTAACTTATGGTGTTCAATGCTTTTCCCGTTATCCGGATCACATGAAACGGCATGACTTTTTCAAGAGTGCCATGCCCGAAGGTTATGTACAGGAACGCACTATATCTTTCAAAGATGACGGGACCTACAAGACGCGTGCTGAAGTCAAGTTTGAAGGTGATACCCTTGTTAATCGTATCGAGTTAAAAGGTATTGATTTTAAAGAAGATGGAAACATTCTCGGACACAAACTCGAGTACAACTTTAACTCACACAATGTATACATCACGGCAGACAAACAAAAGAATGGAATCAAAGCTAACTTCAAAATTCGCCACAACGTTGAAGATGGATCCGTTCAACTAGCAGACCATTATCAACAAAATACTCCAATTGGCGATGGCCCTGTCCTTTTACCAGACAACCATTACCTGTCGACACAATCTAAACTTTCGAAAGATCCCAACGAAAAGCGTGACCACATGGTCCTTCTTGAGTTTGTAACTGCTGCTGGGATTACACATGGCATGGATGAGCTCTACAAATGActagcataaccccttggggcctctaaacgggtcttgaggggttttttg |
| ParR | cccgcgaaattaagatctcgatcccgcgaaattaatacgactcactataggggaattgtgagcggataacaattcccctctagaaataattttgtttaactttaagaaggagatatacataTGATGGACAAGCGCAGAACCATTGCCTTCAAACTAAATCCAGATGTAAATCAAACAGATAAAATTGTTTGTGATACACTGGACAGTATCCCGCAAGGGGAACGAAGCCGCCTTAACCGGGCCGCACTGACGGCAGGTCTGGCCTTATACAGACAAGATCCCCGGACCCCTTTCCTTTTATGTGAGCTGCTGACGAAAGAAACCACATTTTCAGATATCGTGAATATATTGAGATCGCTATTTCCAAAAGAGATGGCCGATTTTAATTCTTCAATAGTCACTCAATCCTCTTCACAACAAGAGCAAAAAAGTGATGAAGAGACCAAAAAAAATGCGATGAAGCTAATAAATTAATTCAATTATTATTGAGTTCCCTTTATCCACTATCAGGCTGGATAAAGGGAACTCAATCAAGTTATTTTCTTACCAGTCATTACATAATCGTTATTATGAAAGGCAGCCATCATCATCATCATCATTAAGgatccggctgctaacaaagcccgaaaggaagctgagttggctgctgccaccgctgagcaataactagcataaccccttggggcctctaaacgggtcttgaggggttttttg |
| AlfB | CCCGCGAAATTAATACGACTCACTATAGGGGAATTGTGAGCGGATAACAATTCCCCTCTAGAAATAATTTTGTTTAACTTTAAGAAGGAGATATACATGTGTCCCGTGAGGACAATTTCATGTATCAAATTAACTGGAATAAGAAAAAATACCCAGAGATTTGCAAGGCACTTGAAGATGCCAAAAATCGTACAGGTGGAATTGCGTGGTATCTTCGTGAGTTGATCCAAAAGGACCTTGAAGAGAAACGCGGTGGTGTAGTCCGTAGCACACCAGTCTACGAGACTGTTGACCAGGAGGTGCAGAACGATCCGCCGAAACCATCCAAGGCTAAGATCGAAACTATCGAGTTGCCTGACGATTCGGGAGGCTTTTTAGAAAATCTGTATTTCCAAAGCCACCACCACCACCACCACTGAGATCCGGCTGCTAACAAAGCCCGAAAGGAAGCTGAGTTGGCTGCTGCCACCGCTGAGCAATAACTAGCATAACCCCTTGGGGCCTCTAAACGGGTCTTGAGGGGTTTTTTG |
| 500 bp Broccoli-Pepper template | cccgcgaaattaatacgactcactatagggAGACCACAACGGTTTccctctagaaataattttgtttaactttaagaaggagatataccaCGGTACCTACCAATCGTGGCGTGTCGGCCTGCTTCGGCAGGCACTGGCGCCGTAGGTACCGGTCGACATCTTTTACTGCAGATGTAACCAGAGAGACACCAAGAGATGAGTCTCCGTATATTTTACTGATCCCCGATAATTTCCCCATTACCTGAGAAATATCTAATGTGGTGCCCCCGAGATCTATAATTAATAAAGAATCTAACTCATCCAGTTCTTGTAGAACTTCATAACCTGCCGGTATAGATTCAGGCATGACTTTTACATCTTTTATTGTGAATGTAGGTGCCTTATTCCGGACGCCGGGCCCGAATGCTGCTACGGCAGTCGAAGACAACATCGCGCCCTTCGGAGGCACCtaaCtagcataaccccttggggcctctaaacgggtcttgaggggttttttg |
| 900 bp Broccoli-Pepper template | cccgcgaaattaatacgactcactatagggAGACCACAACGGTTTccctctagaaataattttgtttaactttaagaaggagatataccaCGGTACCTACCAATCGTGGCGTGTCGGCCTGCTTCGGCAGGCACTGGCGCCGTAGGTACCGGTCGACATCTTTTACTGCAGTTCTGCGTTTTGAAAAAACGTTCATCACGAATCTGTGTGTGTTTTTTTACTGCATCGCATATTAATTCTGCGCCACCGCCTATAACCATAACATGAGTATAACCAGAAAATTCATTGAGCGTATTTAATACACGTTGCTCAAGTTTACGAAGTGCTTCATTCATTGCTTCGGTGACTATTGATATTTTGTTCTCACACCAAGAGATGAGTCTCCGTATATTTTACTGATCCCCGATAATTTCCCCATTACCTGAGAAATATCTAATGTGGTGCCCCCGAGATCTATAATTAATAAAGAATCTAACTCATCCAGTTCTTGTAGAACTTCATAACCTGCCGGTATAGATTCAGGCATGATTTTACATCTTTcTATTGTGAATGTATCCCCGCCATTTAATGTAATTTTTTTCCGGAAGTTTGCTTTCTTACGCTCAATATTTTCCGTATTGGGTTGGTTATTTCTGTCGTAATACTCTGTCAGAGGAAGTGTGCAAACAATATCCACTTCGCTTACCCACTGGTCAGTAAGGCGTGATGCACTGCAACGACATTAACGGGATTCAGGCATGACTTTTACATCTTTGGCAGACTATTGTGAATGTAGGTGCCTTATTCCGGACGCCGGGCCCGAATGCTGCTACGGCAGTCGAAGACAACATCGCGCCCTTCGGAGGCACCtaaCtagcataaccccttggggcctctaaacgggtcttgaggggttttttg |

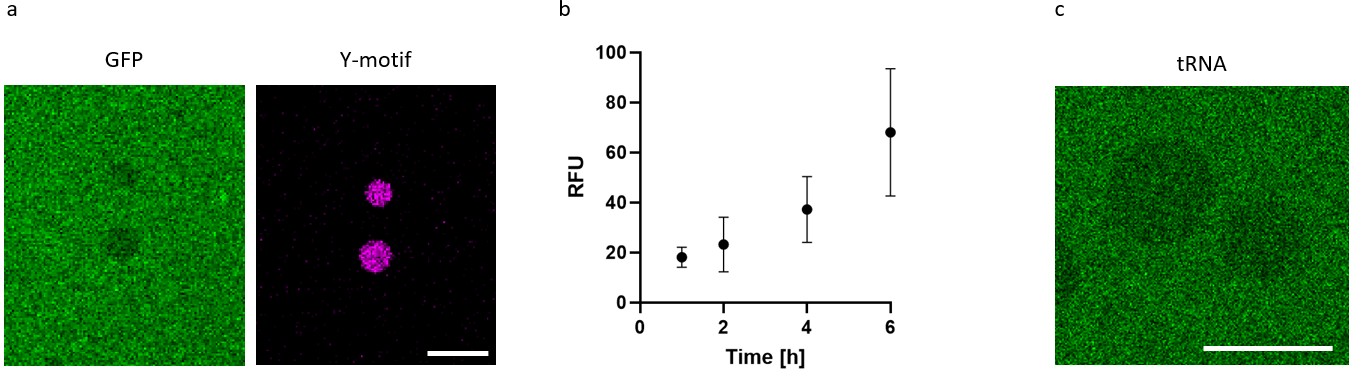

Figure 1: Supplemental material of PURE expression of GFP with Y-motif condensate system. A) Confocal images of product release of GFP (green) into gas phase of condensates (magenta). B) Quantification of relative fluorescence signal of GFP produced in water-in-oil droplets measured at different time points (n = 10/t). C) Confocal image of tRNA loaded with labelled lysine (green) excluded from DNA condensate phase. Scale bars: 10 µm.
